# TAZ (*Wwtr1*) deficiency leads to ER stress and mitochondrial dysfunction in a mouse model of Fuchs’ endothelial corneal dystrophy

**DOI:** 10.64898/2026.02.17.706456

**Authors:** Sangwan Park, Raneesh Ramarapu, Jaegook Lim, Sabina Khan, Meher J. Khan, William R. Stoehr, Brian C. Leonard, Sara M. Thomasy

**Author notes:** SP and RR contributed equally to this work. Address correspondence to: Sara M. Thomasy, Department of Surgical and Radiological Sciences, School of Veterinary Medicine, Department of Ophthalmology & Vision Science, School of Medicine, University of California at Davis, 1 Shields Avenue, 1220 Tupper Hall, Davis, California 95616, USA.

## Abstract

Fuchs’ endothelial corneal dystrophy (FECD) impacts over 300 million individuals worldwide with corneal transplantation as the primary treatment. There is a dire need to establish non-surgical alternatives which are dependent on mouse models. Transcriptional co-activator with PDZ-binding motif (TAZ, encoded by *Wwtr1*) is a mechanotransducer implicated in maintaining homeostasis of corneal endothelial cells (CEnC). *Wwtr1*^-/-^ (TAZ KO) mice serve as an animal model for late-onset FECD. We combined single-cell transcriptomics, transmission electron microscopy, and immunofluorescence staining to elucidate the mechanisms driving pathogenesis in young (2-month-old) and geriatric (11-month-old) mice. A progressive stress response was observed in TAZ KOs defined by endoplasmic reticulum (ER) stress, mitochondrial structural and functional abnormalities, and impaired Na^+^/K^+^ ATPase localization. These changes were accompanied by an altered expression of genes involved in extracellular matrix (ECM) organization, oxidative phosphorylation, macroautophagy and response to oxidative stress. Additionally, we noted age-related differences in cellular response with young TAZ KO CEnCs upregulating macroautophagy and downregulating ECM organization while geriatric TAZ KO CEnCs downregulated macroautophagy, and ECM organization. Both TAZ KO groups downregulated response to oxidative stress and cell-substrate adhesion. Together, these findings establish a mechanistic link between disrupted mechanotransduction and organelle stress in CEnC degeneration, further elaborating on potential mechanisms driving FECD pathogenesis. This positions TAZ KO mice as a translational platform for evaluating non-surgical therapeutic strategies targeting FECD.

**Significance statement:** Fuchs’ endothelial corneal dystrophy (FECD) is a common, age-related cause of vision loss involving a depletion of corneal endothelial cells (CEnC) that necessitates corneal transplantation. Understanding why corneal endothelial cells progressively fail in this disease is essential for developing non-surgical therapies. Using transcriptomics, electron microscopy and immunofluorescence staining, we demonstrate that loss of the mechanotransducer TAZ disrupts cellular homeostasis by inducing endoplasmic reticulum stress, mitochondrial dysfunction and improper extracellular matrix and functional protein organization in CEnCs. By linking altered mechanotransduction to organelle stress and endothelial cell loss, these findings provide insight into fundamental disease mechanisms and identify pathways that may be targeted to preserve corneal endothelial function in FECD.

## Introduction

Corneal endothelial cells (CEnC) are essential to maintain corneal transparency by achieving relative corneal deturgescence through their pump and barrier function. Due to their limited proliferative and regenerative capacity, extensive cell loss and resultant endothelial dysfunction cause persistent corneal edema, often requiring corneal transplantation to restore vision. Fuchs’ endothelial corneal dystrophy (FECD) is a leading cause of corneal endothelial dysfunction involving the formation of guttae or extracellular matrix (ECM) excrescences on Descemet’s membrane (DM) and the gradual loss of CEnCs through apoptosis.

Current hypotheses on the pathogenesis of CEnC apoptosis in FECD include oxidative stress and subsequent mitochondrial and nuclear DNA damage, endoplasmic reticulum (ER) stress or accumulation of unfolded or misfolded proteins in the ER lumen initiating the unfolded protein response (UPR), and abnormal ECM-cell interaction associated with aberrant ECM deposition and TGF-β signaling activating an endothelial-to-mesenchymal transition in CEnCs (1–13). Indeed, growing evidence suggests that multiple mechanisms may eventually converge onto the UPR as a final pathway to mediate CEC apoptosis in FECD (14).

Our group previously investigated the role of the cellular mechanotransducer, transcriptional co-activator with PDZ-binding motif (TAZ), encoded by *Wwtr1* gene in corneal endothelial dysfunction using *Wwtr1*^-/-^ (TAZ KO) mice (15). These mice show progressive decrease in CEnC density, altered zonular occludens 1 (ZO-1) and Na,K-ATPase expression in CEnCs, softer DM, and impaired CEnC regeneration, all of which recapitulate features of FECD patients (15). These characteristics suggest that TAZ KO mice can serve as an animal model for the late-onset FECD. However, there is still a knowledge gap in our understanding of the molecular mechanisms behind how TAZ deficiency leads to CEnC apoptosis and dysfunction.

In this study, we elucidated alterations in CEnC-specific transcriptional profiles from TAZ KO and wildtype (WT) mice at 2 and 11 months of age using single-cell RNA sequencing (scRNA-Seq). We further corroborated our transcriptional data with immunohistochemistry (IHC) and transmission electron microscopy (TEM) of corneal tissue from WT and TAZ KO mice. Our results demonstrate a multifaceted response of CEnCs to TAZ deficiency, highlighting the key role of ER stress and mitochondrial dysfunction. In aggregate, this study provides valuable insight into the dysfunctional mechanisms adopted by CEnCs in FECD, opening avenues to test novel therapeutics by targeting these disease mechanisms using this highly relevant murine model.

## Results

### TAZ KO mice share similar corneal cell type compositions to WT mice

Following quality control, we obtained a total of 29,738 cells across from WT 2-months-old (WT2MO), TAZ KO 2-months-old (KO2MO), WT 11-months-old (WT11MO) and TAZ KO 11-months-old (KO11MO). There were an average of 13,267 transcripts and 2891 unique genes per cell. Following batch correction via *Harmony*, unsupervised clustering revealed the presence of 9 clusters (16). Quality control plots for each cluster can be found in **Supplementary File 1S2**. We identified 8 major cell types including limbal stem cells (LSC) (cluster 5), basal cells (BC) (cluster 0 and 2), wing cells (WC) (cluster 1), superficial cells (SC) (cluster 3), conjunctival epithelium (CjE) (cluster 4), CEnC (cluster 6), dendritic cells (DC) (cluster 7) and stromal keratocytes (CSK) (cluster 8) in all 4 samples (**Fig 1A and B**). Clusters are numbered by size and no cell clusters were specific to any sample (**Fig 1A-C**). We identified a distinct cluster of CEnC in all 4 samples based upon the expression pattern of known markers (*Col8a2, Col8a1, Col4a2, Slc4a11, Car3* and *Atp1b1*) (**Fig 1A and B**) (17,18). Consistent with their genotypes, we identified no *Wwtr1* transcripts in either KO2MO or KO11MO samples (**Fig 1D**). Further analysis was performed within the CEnC cluster to investigate the transcriptomic alterations of CEnC between ages and genotypes.

**Figure 1.**
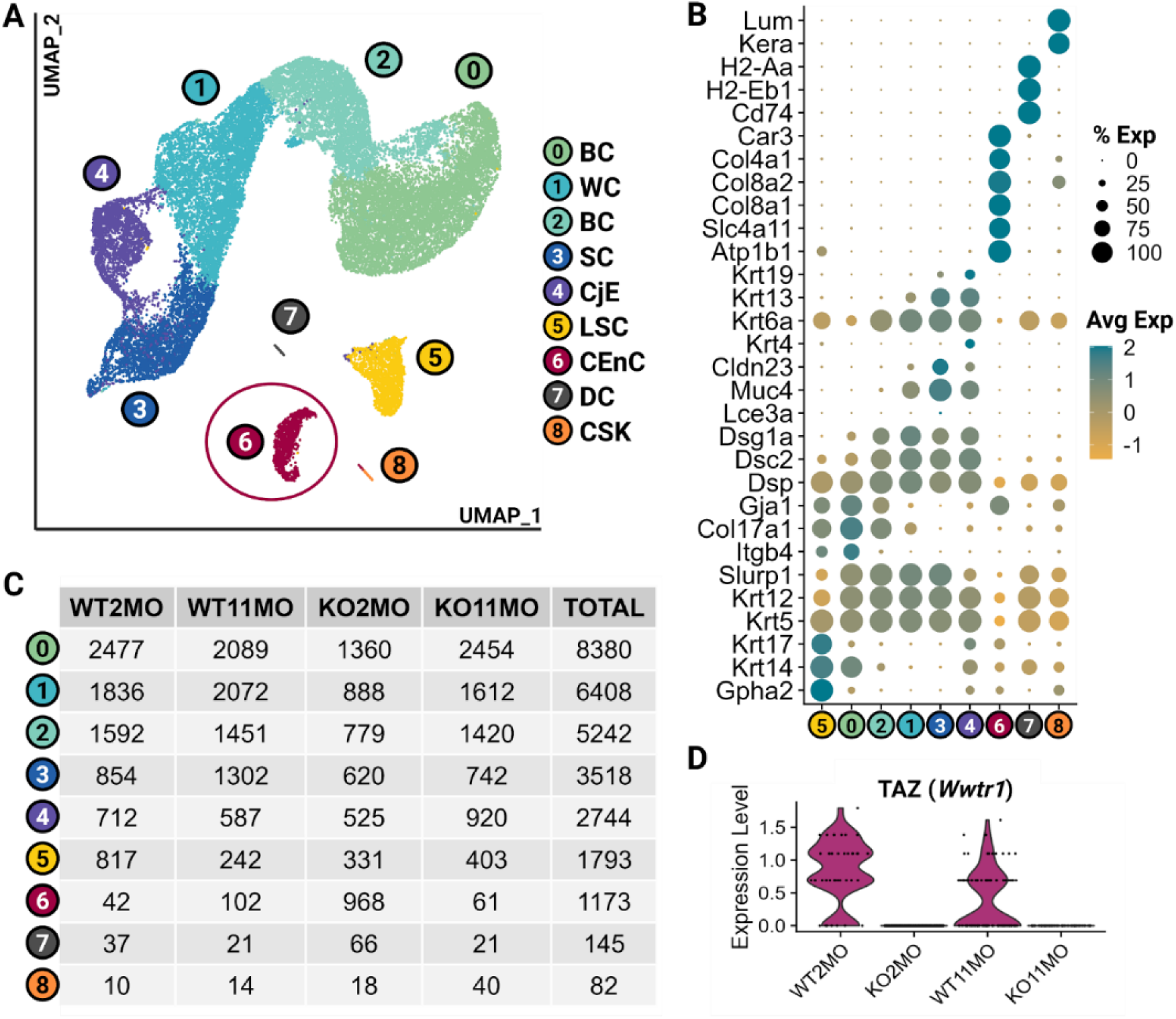
The scRNA-seq atlas of corneal cells in wildtype (WT) and *Wwtr1*^-/-^ (TAZ KO) mice reveals 8 clusters including corneal endothelial cells (CEnC<u>s</u>). (A) UMAP demonstrating the unsupervised clustering results for murine corneal cells from WT and TAZ KO at 2 and 11 months of age. (B) Dot plot demonstrating cell type specific marker expression of the 8 clusters. (C) Table of cell counts for each cluster by sample. (D) Violin plot of *Wwtr1* expression in CEnC (cluster 6) by sample. Limbal stem cells, LSC; basal cells, BC; wing cells, WC; superficial cells, SC; conjunctival epithelium, CjE;; dendritic cells, DC; corneal keratocytes, CSK.

### Young TAZ KO mice showed upregulation of macroautophagy and downregulation of aerobic respiration and ECM synthesis

We performed differentially expressed gene (DEG) analysis to investigate the transcriptomic differences between WT and TAZ KO CEnC at 2 months of age (WT2MO and KO2MO, respectively) via Wilcoxon-Rank-Sum testing. The DEG analysis of WT2MO and KO2MO produced 2,917 DEG of the total 12,276 genes assessed (23.8%). The Gene ontology (GO) over-enrichment analysis (ORA) between genotypes at 2 months of age revealed an enrichment of multiple biological processes (**Supplementary File 2**). The top depleted GO terms in KO2MO versus WT2MO included genes involved in aerobic respiration specifically mitochondrial respiratory chain complexes (e.g. *Cox5a*, *Cox7a*, *Atp5b*, *Ndufa7*) and ECM synthesis/organization (e.g. *Col1a1*, *Col3a1*, *Vim*) as shown in **Fig 2A and B** and **Supplementary File 1S3**. The top enriched GO terms in KO2MO versus WT2MO included genes involved in regulation of apoptotic signaling pathway, particularly *Bnip3*-mediated mitophagy process (e.g. *Bnip3* and *Bnip3l*) and *Wnt* signaling (*Wnt6*, *Fzd1*, *Ctnnd1*, *Lrp5*) as detailed in **Fig 2B and C** and **Supplementary File 1S3**. Additionally, we noted a depletion of “negative regulation of apoptotic signaling” in KO2MO compared to WT2MO suggesting a constitutive anti-apoptotic process is disrupted in the KO2MO.

**Figure 2.**
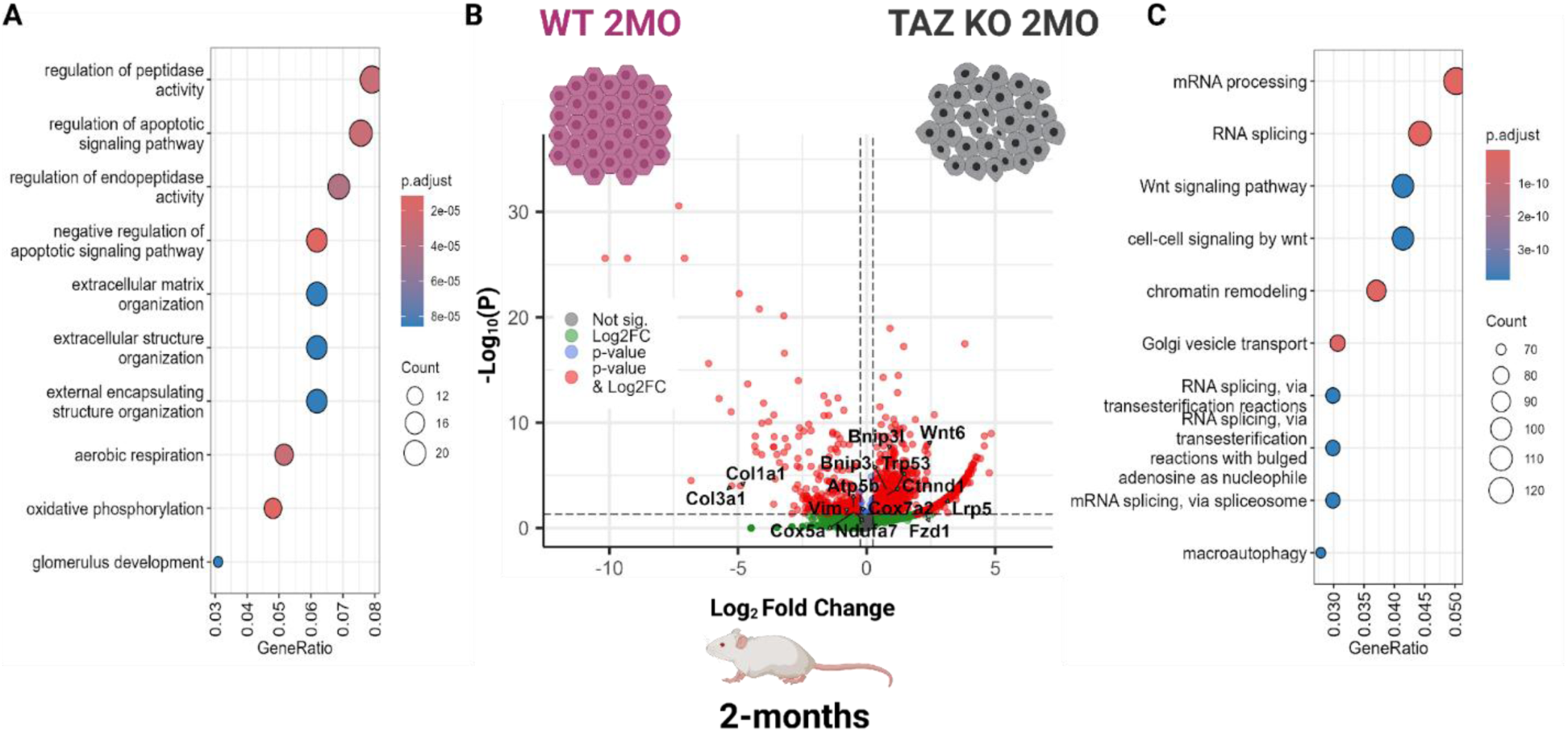
Young *Wwtr1*^-/-^ (TAZ KO) mice upregulate macroautophagy but downregulate aerobic respiration and extracellular matrix synthesis in the corneal endothelium. (A) Dot plot demonstrating over-representation gene ontology (biological process) for differentially expressed genes (DEG) upregulated in WT compared to TAZ KO at 2 months of age (WT2MO and KO2MO, respectively). (B) Volcano plot of DEG expression results between WT2MO and KO2MO (Wilcoxon-Rank-Sum). (C) Dot plot demonstrating over-representation gene ontology (biological process) for DEG upregulated in KO2MO compared to WT2MO. *Wwtr1* was removed from this visualization due to its outlier status.

### Old TAZ KO mice showed downregulation of macroautophagy, mitochondrial respiratory chain complexes and ATP production

We additionally performed DEG analysis and GO ORA between WT and TAZ KO CEnC at 11 months of age (WT11MO and KO11MO, respectively). The DEG analysis of WT11MO and KO11MO produced 2178 DEG of the total 12,276 genes assessed (17.7%). As found in **Supplementary File 3**, the top depleted GO terms for KO11MO compared to WT11MO included genes involved in autophagy, particularly autophagosome-related macroautophagy (e.g. *Atg10*, *Atg13*, *Rab1b*), and biogenesis and metabolism of various nucleoside triphosphates including ATP (e.g. *Atp5g3, Atpsckmt, Atp6*). Additionally, downregulation of genes involved in aerobic respiration (*Cox4l2*, *Ulk2*) and mitochondrial respiratory chain complexes (e.g. *mt-Co1*, *mt-Co2*, *mt-Nd1*, *mt-Nd4*, *mt-Nd6*) were identified in KO11MO versus WT11MO (**Fig 3A and B; Supplementary File 1 S4**). The top enriched GO terms in KO11MO compared to WT11MO were mostly associated with RNA and protein processing, but there were substantially fewer upregulated genes in KO11MO (**Figure 3B and C; Supplementary File 1 S4**).

**Figure 3.**
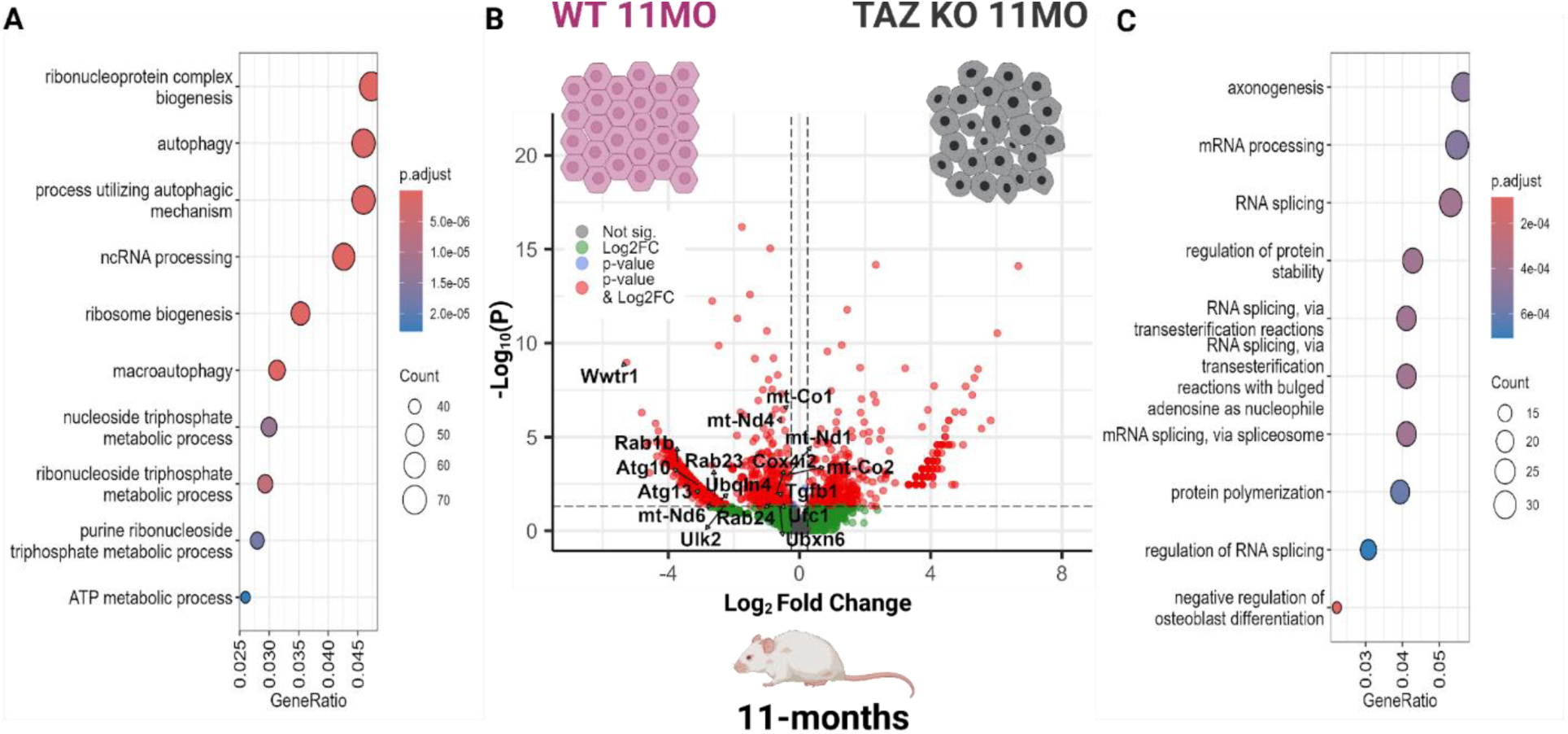
Old *Wwtr1*^-/-^ (TAZ KO) mice downregulate macroautophagy and aerobic respiration in corneal endothelium. (A) Dot plot demonstrating over-representation gene ontology (biological process) for differentially expressed genes (DEG) upregulated in WT compared to TAZ KO at 11 months of age (WT11MO and KO11MO, respectively). (B) Volcano plot of DEG expression results between WT11MO and KO11MO (Wilcoxon-Rank-Sum). (C) Dot plot demonstrating over-representation gene ontology (biological process) for DEG upregulated in KO11MO compared to WT11MO.

### Key pathways implicated in FECD are altered in TAZ KO mice independent of age

In order to understand the age-independent effect of *Wwtr1* deficiency on the CEnC transcriptomic landscape, we identified the shared upregulated and downregulated genes across both ages (**Figure 4A**). We found 272 shared upregulated and 84 shared downregulated genes in CEnCs of TAZ KO mice of both age groups compared to their age-matched WT groups (**Figure 4A; Supplementary File 4**). The GO ORA for biological process demonstrated an enrichment of the GO terms in TAZ KO CEnCs including cellular component disassembly (e.g. *Ccn2, Dedd2, Ltn1*), mesenchyme development (*e.g. Fgf9, Foxc1, Nog, Pitx2, Sox4*) and cell cycle G1/S phase transition (e.g. *Cables1, Ccnd2, Cdk2ap2, Rfwd3, Tbx2*) (**Fig 4B and D-G; Supplementary File 1 S5; Supplementary File 4**). By contrast, response to oxidative stress (e.g. *Hif1a, Hmox1, Sod3*, *Car3*), cell-substrate adhesion (*Npnt, Nid1, Spock1*), and ECM organization (*Col4a1, Col1a1, Fbln5*) were shared downregulated GO terms in the TAZ KO CEnCs (**Fig 4C and H-K; Supplementary File 1 S5; Supplementary File 4**).

**Figure 4.**
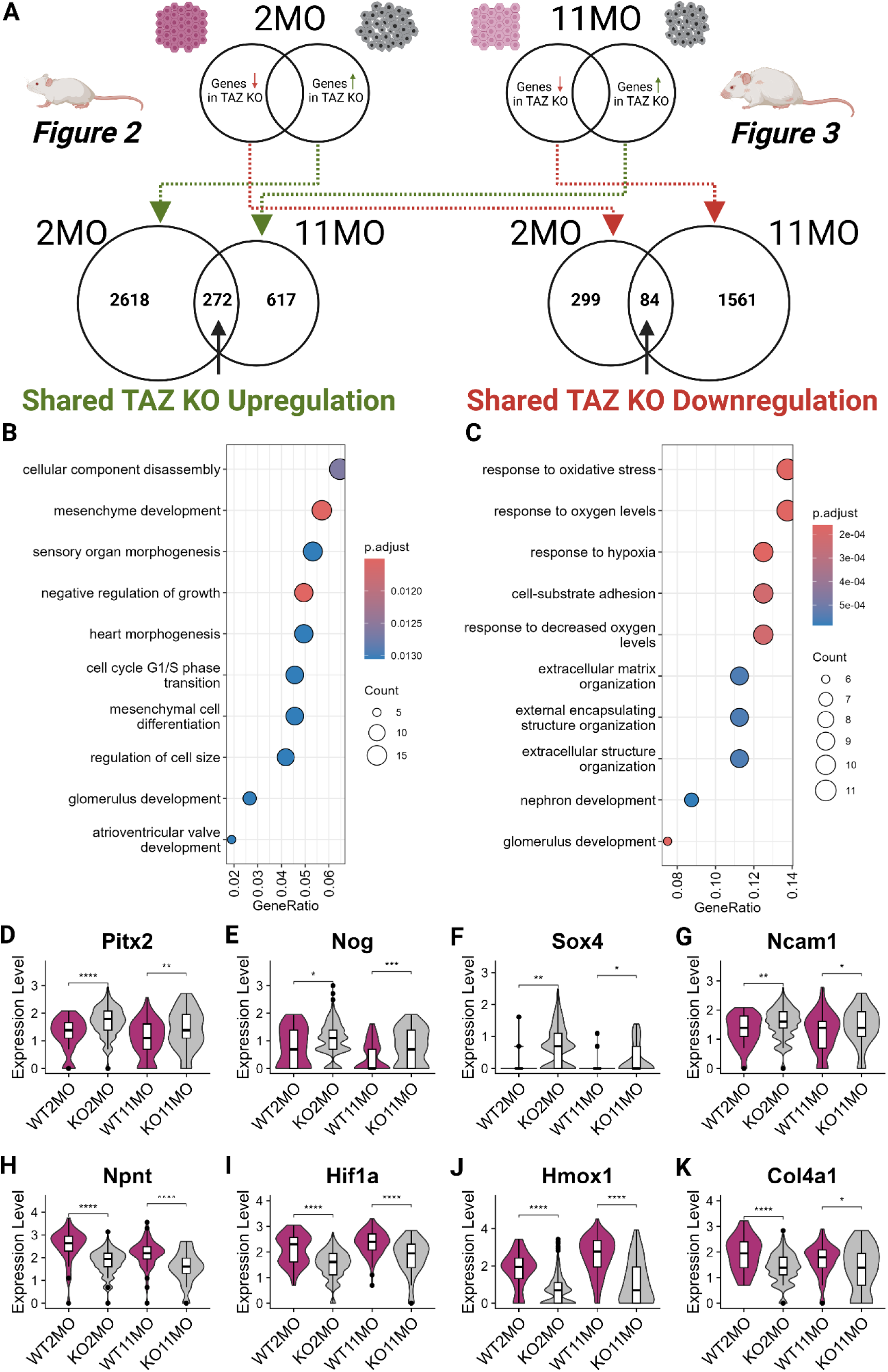
Key pathways implicated in FECD are altered in corneal endothelial cells (CEnCs) of *Wwtr1*^-/-^ (TAZ KO) mice independent of age. (A) Flowchart illustrating identification of the shared differentially expressed genes (DEG) set. (B) Dot plot demonstrating over-representation gene ontology (biological process) for shared DEG upregulated by TAZ KO CEnC. (C) Dot plot demonstrating over-representation gene ontology (biological process) for shared DEG downregulated by TAZ KO CEnC. (D-G) Violin plots of select genes from GO ORA upregulated in TAZ KO compared to WT CEnC independent of age. (H-K) Violin plots of select genes from GO ORA downregulated in TAZ KO compared to WT CEnC independent of age.

### TAZ deficiency leads to ER stress, mitochondrial dysfunction and resultant CEnC loss

Corneal transmission electron microscopy (TEM) images from TAZ KO mice revealed marked abnormalities in DM and CEnC morphology. TAZ KO mice exhibited a thinner DM when compared with WT mice (**Fig 5**), consistent with our previous study whereby TAZ KO mice had an overall thinner cornea (15). Additionally, this is consistent with the depletion of genes associated with DM such as *Col4a1, Col1a1*, and *Col3a1* in the TAZ KO CEnC transcriptome (**Fig 2B, 4C and K; Supplementary File 1 S5; Supplementary File 4**).

**Figure 5.**
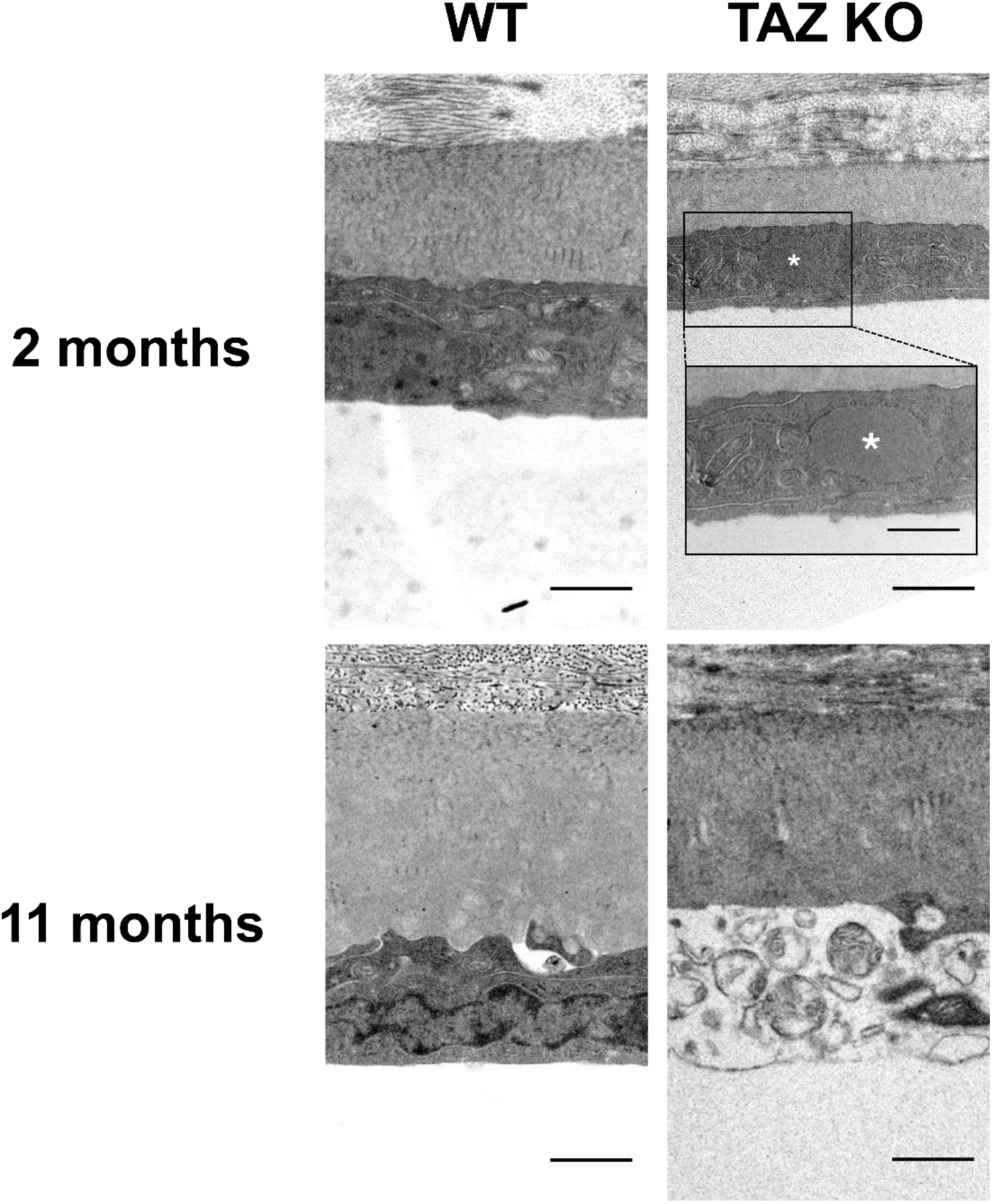
*Wwtr1*^-/-^ (TAZ KO) mice showed ultrastructural evidence of endoplasmic reticulum (ER) stress and resultant death of corneal endothelial cells (CEnCs). Transmission electron microscopy revealed that TAZ KO mice showed dilated ER (asterisk) starting at 2 months of age and perinuclear electrolucency and extensive vacuolation with engulfed and digested cytoplasmic organelles as they age, eventually leaving bare Descemet’s membrane without CEnC in some regions. Scale bar = 1 μm (500 nm in the inset).

Additionally, CEnCs from TAZ KO mice showed dilated ER (**Fig 5 inset**) and abnormal appearance of mitochondria characterized as indistinct cristae and electrolucency in the inner mitochondria membrane and intermembrane space beginning at 2 months of age (**Fig 6A**). By 12 months of age, the severity had increased with numerous autophagosomes and regional CEnC loss also identified (**Fig 5**). Although ER stress was not significantly different at the transcriptional level, the known ER stress marker protein, GRP78 (78 kDa glucose-regulated protein) was more highly expressed in CEnCs of TAZ KO mice versus WTs (**Fig 7**). Additionally, TAZ KO mice showed nuclear translocation of GRP78, which was not observed in WT (**Fig 7A, white arrows**).

**Figure 6.**
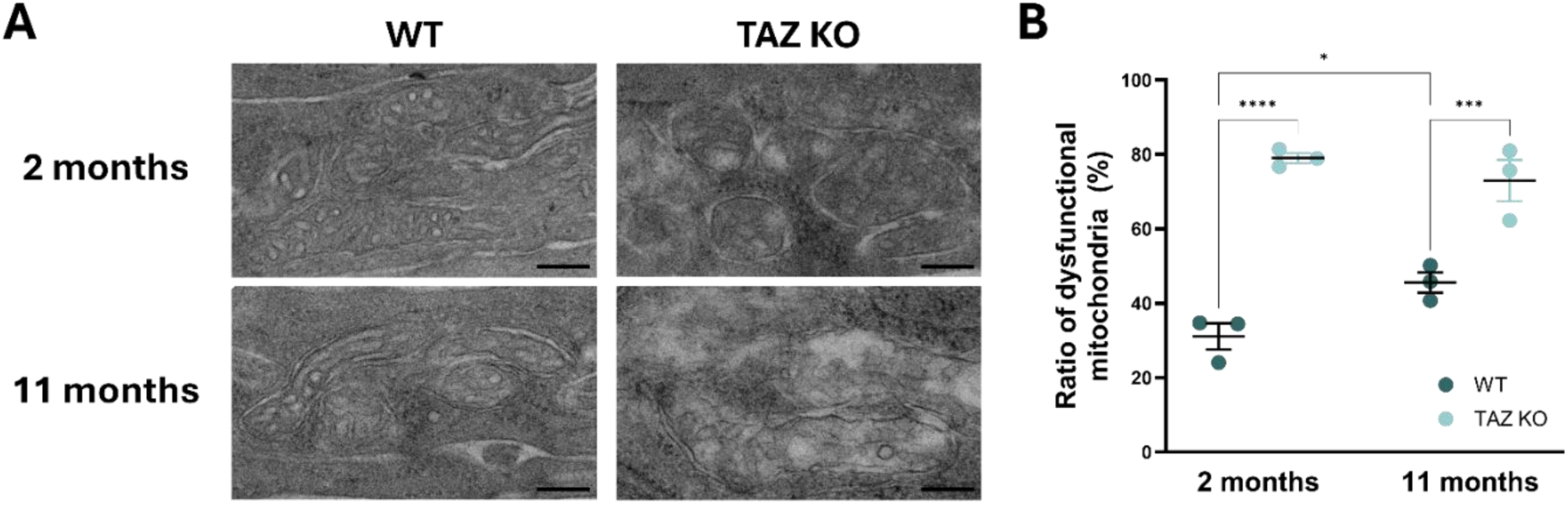
*Wwtr1*^-/-^ (TAZ KO) mice showed more abundant dysfunctional mitochondria and downregulation of mitochondrial genes in corneal endothelium compared to wildtype (WT) mice. (A) TAZ KO mice showed indistinct cristae and electrolucency in the inner mitochondria membrane and intermembrane space at 2 months of age with an increase in severity by 11 months of age (scale bar = 200 nm). (B) The ratio of dysfunctional mitochondria was significantly greater in TAZ KO versus WT mice within each age group (2-way ANOVA).

**Figure 7.**
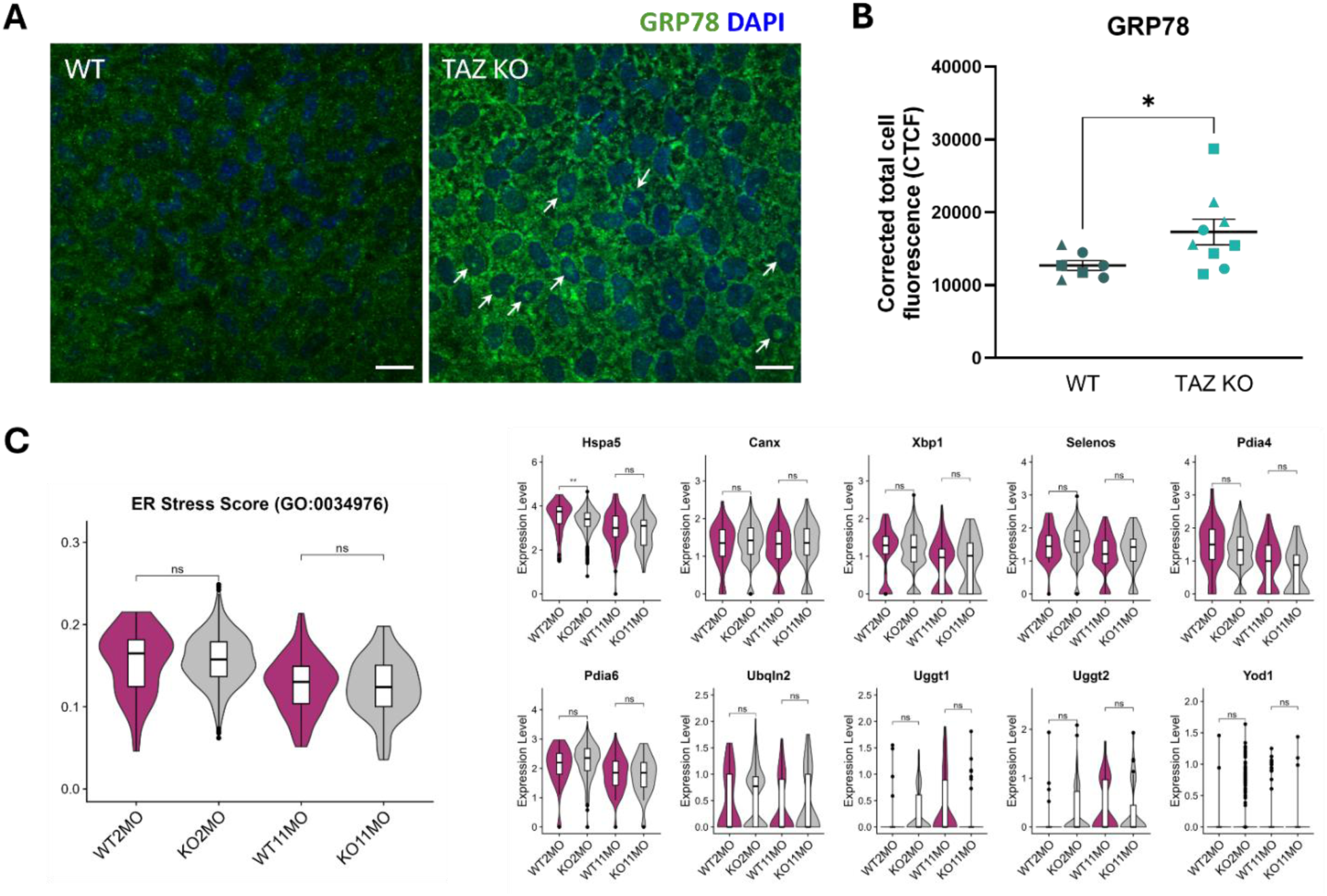
*Wwtr1*^-/-^ (TAZ KO) mice showed overexpression of GRP78 in corneal endothelium, suggesting that ER stress is involved in the pathogenesis. (A) Immunofluorescence staining of GRP78 in corneal endothelial wholemounts from wildtype (WT) and TAZ KO mice (scale bar = 20 µm). The arrow indicates nuclear translocation of GRP78. (B) TAZ KO mice showed significantly greater GRP78 expression in corneal endothelium compared to WT mice (unpaired t-test). Triangles, ≤ 2 months of age; rectangles, > 10 months of age; circles: 3-10 months of age. (C) Violin plots of ER stress-related genes in WT and TAZ KO CEnC at 2 and 11 months of age.

The ratio of dysfunctional mitochondria was significantly greater in TAZ KO versus WT mice in both 2- and 12-month age groups (**Fig 6B**). From a transcriptional standpoint, we identified no significant differences in percentage mitochondrial gene expression score between the WT and TAZ KO CEnC at 2 months of age; however, we noted a significant decrease in percentage mitochondrial genes in the TAZ KO CEnC at 11 months of age compared to the WT (**Fig 8**). These differences were driven by select mitochondrial genes: *mt-Nd1, mt-Nd4, mt-Nd5, mt-Co1, mt-Co2, mt-Co3, mt-Cytb and mt-Atp6* (**Fig 8 and Supplementary File 1 S6**). It is important to note that mitochondrial genes showed an overall higher expression at 11 months of age compared to 2 months independent of genotype, suggesting that gene expression differences are only found later as the disease progressed and fewer cells are present (**Supplementary File 1 S6**). Additionally, the staining intensity of ATP synthase, a key component of aerobic respiration in mitochondria, was significantly lower in CEnC of TAZ KO versus WT mice (**Fig 8**).

**Figure 8.**
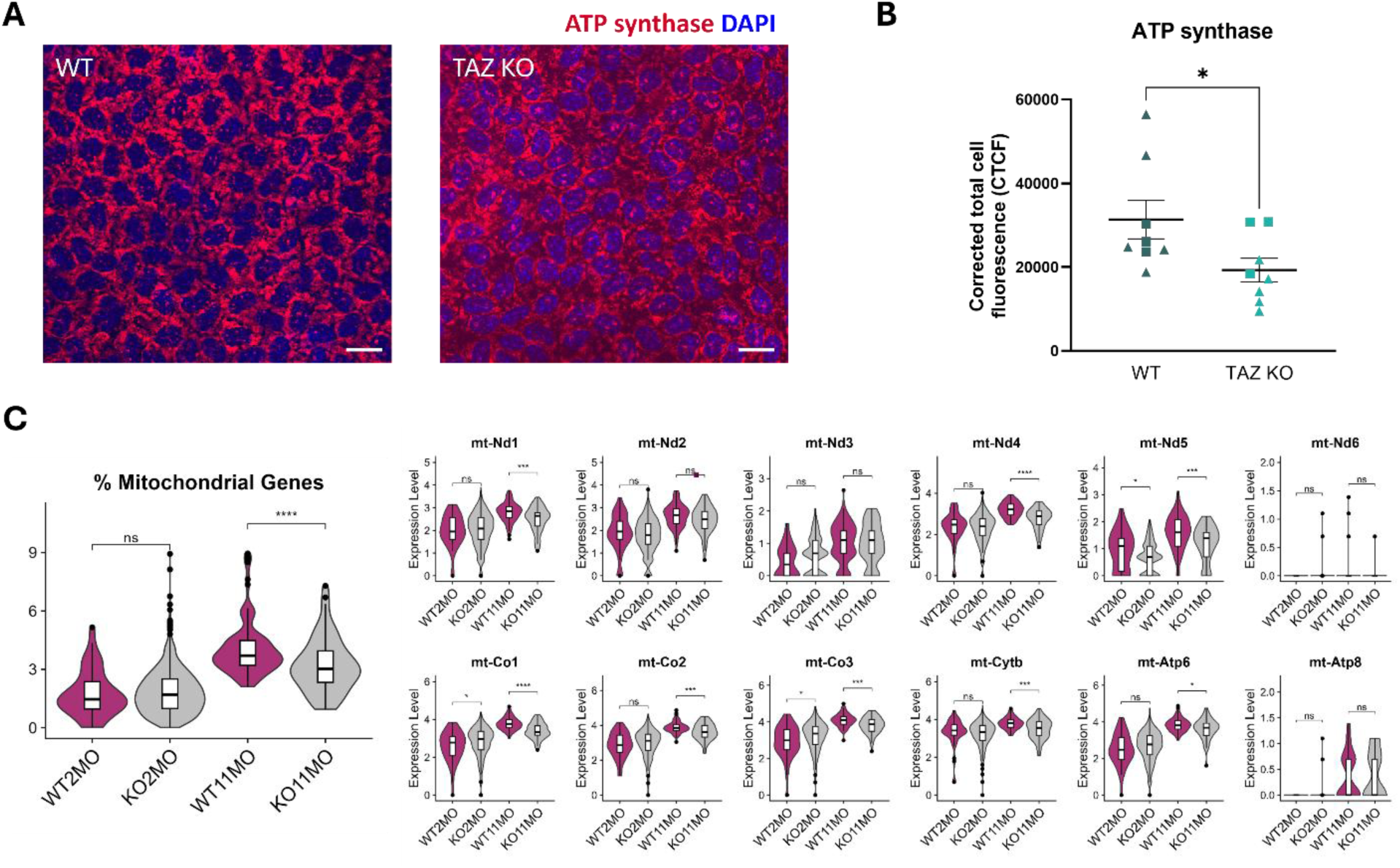
*Wwtr1*^-/-^ (TAZ KO) mice showed reduced expression of ATP synthase and downregulation of mitochondrial genes in corneal endothelium, suggesting that mitochondrial dysfunction plays a role in the pathogenesis. (A) Immunofluorescence staining of ATP synthase in corneal endothelial wholemounts (scale bar = 20 µm). (B) TAZ KO mice showed significantly lower ATP synthase expression in corneal endothelium compared to wildtype (WT) mice (unpaired t-test). Triangles, ≤ 2 months of age; rectangles, > 10 months of age. (C) Violin plots of mitochondrial genes in WT and TAZ KO CEnC at 2 and 11 months of age.

### Aberrant localization of ATP1A1 and upregulation of certain pump and barrier function-related transcripts in TAZ KO CEnC

The expression of ATP1A1 (α-1 subunit of Na,K-ATPase), the pump function marker of CEnCs, appeared altered morphologically in TAZ KO mice at 12 months of age. While some regions showed diminished and discontinuous expression of ATP1A1 along the intercellular border in old TAZ KO mice, the remainder of CEnCs of old TAZ KO mice showed more diffuse localization along the intercellular border compared to WT mice (**Fig 9A**). This aberrant localization of ATP1A1 on CEnC wholemounts was found with a downregulation of its respective gene, *Atp1a1*, in TAZ KO mice. Interestingly, we noted unique changes at the transcriptional level to the expression of other subunits of Na,K-ATPase. We identified an upregulation of *Atp1b1* and *Atp1b3* genes encoding the β-subunits of Na,K-ATPase in CEnC of TAZ KO mice (**Fig 9B**).

**Figure 9.**
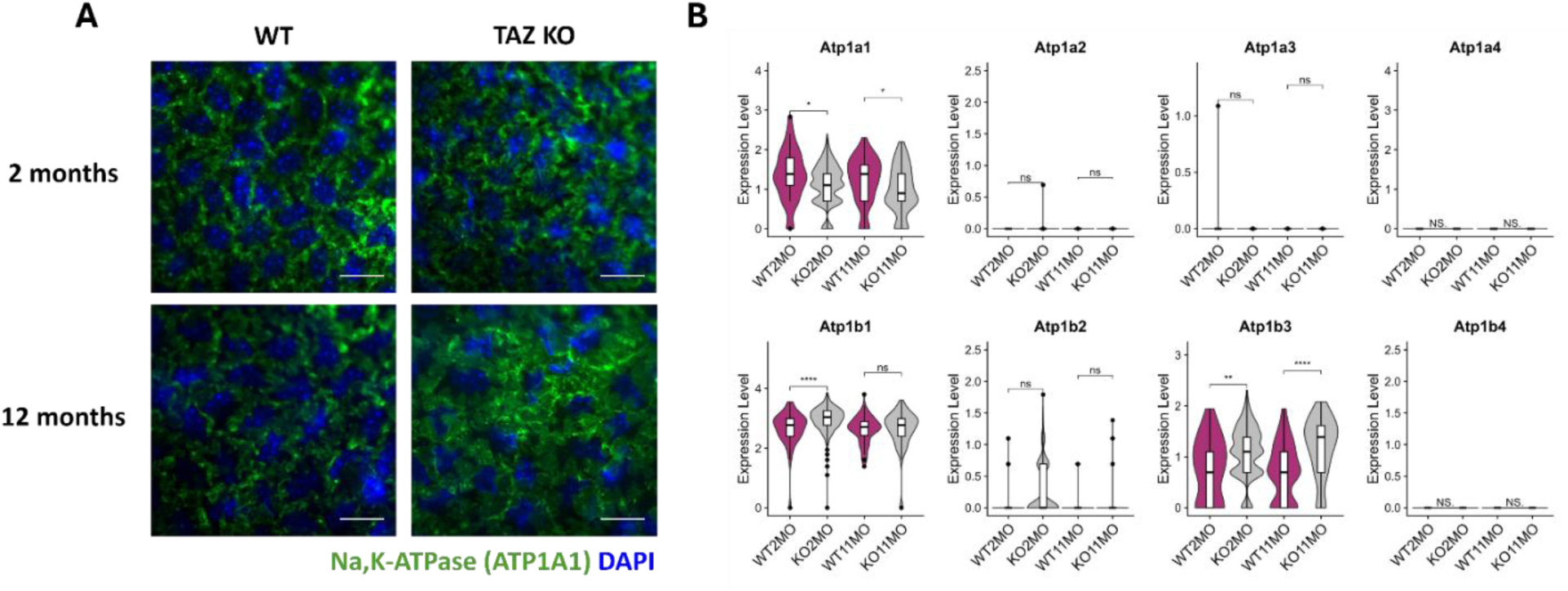
*Wwtr1*^-/-^ (TAZ KO) mice showed aberrant localization of ATP1A1 (α-1 subunit of Na,K-ATPase) and dysregulated expression of genes related to Na,K-ATPase subunits in corneal endothelial cells (CEnC). (A) Immunofluorescence staining of ATP1A1 in corneal endothelial wholemounts from wildtype (WT) and TAZ KO mice (scale bar = 20 µm). Adapted from “Mice Deficient in TAZ (Wwtr1) Demonstrate Clinical Features of Late-Onset Fuchs’ Endothelial Corneal Dystrophy,” by BC Leonard, S Park, S Kim et al, 2023. Invest Ophthalmol Vis Sci, 64(4):22 (http://doi:10.1167/iovs.64.4.22). CC BY-NC-ND 4.0. (B) Violin plots of genes related to Na,K-ATPase subunits in WT and TAZ KO mice CEnC at 2 and 11 months of age.

Although ZO-1 expression was regionally disrupted in TAZ KO mice from 2 months of age in our previous study,^15^ its respective gene, *Tjp1* showed no significant difference at the transcriptional level (**Fig 10**). However, other critical junctional proteins *Cdh2* and *Ncam1* were significantly upregulated in CEnC of TAZ KO mice. Particularly, *Cdh2*, one of the demonstrated endothelial markers in mice, was significantly upregulated in CEnC of TAZ KO mice versus WT at 11 months of age (**Fig 10**).

**Figure 10.**
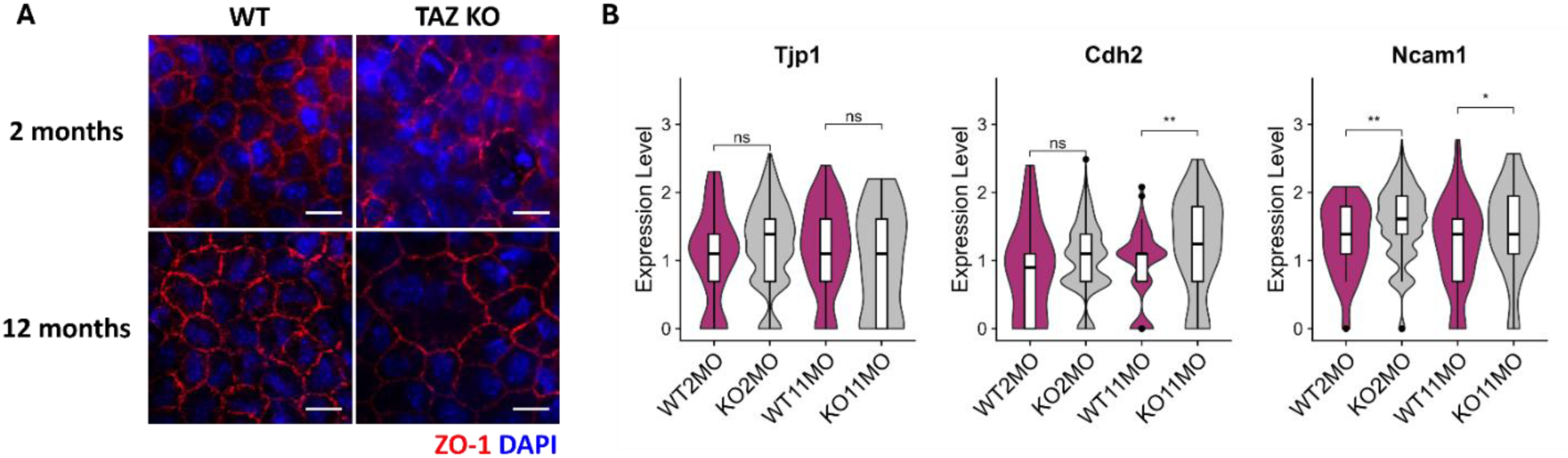
*Wwtr1*^-/-^ (TAZ KO) mice showed upregulation of some junctional marker genes (*Cdh2* and *Ncam1*) of corneal endothelial cells. (A) Immunofluorescence staining of ZO-1 in corneal endothelial wholemounts from wildtype (WT) and TAZ KO mice (scale bar = 20 µm). Adapted from “Mice Deficient in TAZ (Wwtr1) Demonstrate Clinical Features of Late-Onset Fuchs’ Endothelial Corneal Dystrophy,” by BC Leonard, S Park, S Kim et al, 2023. Invest Ophthalmol Vis Sci, 64(4):22 (http://doi:10.1167/iovs.64.4.22). CC BY-NC-ND 4.0. (B) The expression level of *Tjp1* gene encoding ZO-1 was not significantly different between genotypes. *Cdh2* was upregulated in TAZ KO mice at 11 months of age versus WT mice. *Ncam1* was regulated in TAZ KO mice at both 2 and 11 months of age versus WT mice.

## Discussion

In the current study, our results demonstrate that the gradual CEnC loss in TAZ KO mice involves ER stress and mitochondrial dysfunction. Although our scRNA-sequencing showed no transcriptional changes in *Hspa5* (encodes GRP78, a molecular chaperone in the ER), we identified that GRP78 was more highly expressed and translocated into the nucleus of CEnCs in TAZ KO mice. Indeed, ER stress is known to cause nuclear translocation of GRP78 in normal lung epithelial cells (19). In models of chronic ER stress, cells initiate a feedback mechanism that actively suppresses the transcription of certain mRNAs, including *Hspa5*, possibly through pathways like IRE1-dependent decay (RIDD) and the silencing of ATF6α (20). In this scenario, the elevated protein amounts are maintained primarily through post-transcriptional mechanisms, and an upregulation of mRNA is no longer the main driver of increased GRP78 expression (21,22). Overexpression of GRP78 indicates that cells have initiated protective mechanisms against cell death caused by disturbances of ER homeostasis and if this ER stress persists despite GRP78 overexpression, apoptosis is subsequently triggered (23,24). Consistent with this study, increased GRP78 expression was demonstrated in human corneas affected with FECD, cultured FECD-affected CEnCs and *Col8a2* (L450W/L450W) knock-in mice (6,8–11,25). In addition to the increased production of ER chaperones and protein-folding enzymes in the ER, the UPR also could increase the size of the ER to dilute the unfolded protein load. Indeed, TAZ KO mice showed dilated ER starting at 2 months of age.

Additionally, TAZ KO mice manifest ultrastructural evidence of mitochondrial pathology along with significant downregulation of mitochondrial genes beginning at 2 months of age. Most of these DEGs including *mt-Nd1*, *mt-Nd4*, and *mt-Nd5* encode nicotine-amide adenine dinucleotide (NADH) dehydrogenase, which is critical for the electron transport in the mitochondrial respiratory chain complex I. The genes encoding mitochondrial cytochrome c oxidase (*mt-Co1*, *mt-Co2*, and *mt-Co3*) and the *mt-Cytb* gene encoding mitochondrial cytochrome b in the respiratory chain complex IV and III, respectively, were also downregulated in TAZ KO mice. Additionally, the genes encoding the subunits of the respiratory chain complex IV (*Cox4l2, Cox5a*, and *Cox7a*) were also downregulated. Finally, the *Atp5* and *mt-Atp6* genes encoding adenosine triphosphate (ATP) synthase in complex V, the final enzyme in mitochondrial oxidative phosphorylation to generate ATP, was also downregulated in TAZ KO mice versus WT mice. In addition to the transcriptomic evidence, IHC of ATP synthase shows a significant downregulation in TAZ KO compared to WT which is further suggestive of dysregulation. Furthermore, the fact that the electron transport chain and ATP synthase are embedded in the mitochondrial inner membrane where cristae are formed corroborates the localization of mitochondrial morphological changes observed on TEM (26). These abnormal mitochondrial bioenergetics identified in TAZ KO mice recapitulate the known pathophysiology of FECD, which includes loss of mitochondrial membrane potential, reduced ATP synthesis by decreased cytochrome oxidase activity or absence of complex I and V enzymes in the mitochondria electron transport chain, and increased production of mitochondrial ROS (27–30).

Interestingly, TAZ KO mice at 2 months of age showed upregulation of *Bnip3* and *Bnip3l*, which may indicate the initiation of *Bnip3*-mediated mitophagy process in CEnCs. Mitophagy is a type of macroautophagy that eliminates damaged mitochondria in order to maintain mitochondrial homeostasis. *Bnip3*-mediated mitophagy is triggered by mitochondrial membrane depolarization, hypoxia, or post-translational modifications and mediated by receptor proteins on the outer mitochondrial membrane such as BNIP3 binding directly to the LC3 on the autophagosome membranes (31,32) Excessive mitophagy, however, leads to over-clearance of mitochondria and insufficient cellular energy supply, resulting in cell death (31,32). In TAZ KO mice, the number of dysfunctional mitochondria increased with age, eventually leading to extensive digestion of intracellular organelles and regional CEnC loss at 11 months of age.

Mitochondria are the primary source of ATP that are necessary for an active pump function of Na,K-ATPase to keep the cornea dehydrated (33). Clinically, TAZ KO mice do not develop corneal edema but showed upregulation of the β-subunits of Na,K-ATPase (*Atp1b1* and *Atp1b3*) as well as the junctional markers including *Cdh2* and *Ncam1* (15). The upregulation of certain transcripts for pump and barrier functions may suggest a compensatory response of the remaining CEnCs to maintain corneal deturgescence. Alternatively, it may be a part of the disease process in TAZ KO mice. There have been reports suggesting a relationship between ER stress and assembly of the Na,K-ATPase subunits in the ER. Na,K-ATPase proteins undergo post-translational maturation in the ER prior to exportation to the Golgi and plasma membrane and GRP78 plays a crucial role in the folding of the β-subunit of Na,K-ATPase (34). Na,K-ATPase β-subunits serve as a chaperone for the α-subunit (34). Overexpression of Na,K-ATPase β-subunits resulted in upregulation of the α-subunits and increased the membrane-bound Na,K-ATPase abundance improving pulmonary edema in a rat model of congestive heart failure (35,36). In TAZ KO mice, however, upregulation of β-subunits was associated with downregulation of α-subunits at the transcriptional level. It was demonstrated that individual subunits cannot exit the ER and only the Na,K-ATPase α-β complexes can be exported to the Golgi and the number of α_1_ subunits determines the amount of the α-β complexes (34,37). The aberrant localization of the α_1_-subunit of Na,K-ATPase protein demonstrated in TAZ KO mice in conjunction with the downregulation of α-subunits at the transcriptional level suggests a potential inherent defect in maturation of Na,K-ATPase in CEnC of TAZ KO mice.

Given that aberrant localization of ATP1A1 expression was not observed in young TAZ KO mice, we presume pre-existing ER stress would be an exacerbating factor for maturation of Na,K-ATPase in CEnC of TAZ KO mice. The chronological relationship between the maturation of Na,K-ATPase and ER stress, however, has yet to be determined. Furthermore, it is unclear whether ER stress preceded mitochondrial dysfunction in TAZ KO mice as evidence of ER stress and morphological alteration of mitochondria occurred concomitantly at 2 months of age. Challenging the relationship of increased ER stress to mitochondrial dysfunction with various ER stress inhibitors could clarify the pathogenic role of ER stress in TAZ-deficient mice. For example, blocking GRP78 activation with systemic N-acetyl-L-cysteine (NAC) increased CEnC survival in the Col8a2 mutant mice (25). In a diabetic CEnC dysfunction mouse model, injecting the ER stress antagonist 4-phenylbutyric acid (4-PBA) mitigated corneal edema development via improving mitochondrial bioenergetic deficiency (38). It is also possible that ER stress and mitochondrial dysfunction reinforce each other and propagate a vicious cycle, which leads to apoptosis. Calcium is considered a key communicator between the ER and mitochondria as Ca^2+^ transfer from the ER to mitochondria is required for initiation of apoptosis and mitochondrial Ca^2+^ overload results in decreased ATP production and increased ROS generation (39).

Finally, a notable finding from our transcriptional data reveals the activation of a potential regenerative mechanism in our mouse model in response to stress. Among the shared upregulated genes independent of age, we identified expression of early CEnC genes involved with the differentiation of its precursor neural crest-derived periocular mesenchyme. These include transcription factors such as *Pitx2*, *Foxc1* and *Wnt* signaling members. Specifically, *Pitx2* functions as a negative regulator of *Foxc1*, influencing a common pathway and allows for integral fine tuning of *Foxc1* dosage (40,41). Together with *Wnt* activation, these factors drive periocular mesenchyme multipotency and differentiation into CEnC (41,42). Coupled with an enrichment for G1/S phase transition (proliferative activation) and a downregulation of terminal development CEnC adhesion markers (*Nid1* and *Npnt*), our data suggests a regenerative attempt by TAZ KO CEnC (43). It is prudent to consider this within the context of the mouse model given the known differences in regenerative capacities between the mouse and human CEnC (44). Nonetheless, this highlights model caveats and opens an avenue towards considering potential regenerative therapeutics.

In conclusion, we demonstrated that ER stress and mitochondrial dysfunction drive CEnCs to a pathological state with impaired endothelial pump function and regional CEnC loss in TAZ KO mice. This highlights that *Wwtr1* plays a critical role in unleashing the cell death mechanism in CEnCs via ER stress and mitochondrial dysfunction.

## Materials and Methods

### Animals

Strain 129S WT and TAZ KO mice were included in this study. We adhered to all guidelines from the Association for Research in Vision and Ophthalmology Statement for the Use of Animals in Ophthalmic and Vision Research and this study was performed in accordance with the National Institutes of Health (NIH) Guide for the Care and Use of Laboratory Animals. We performed dissections according to a protocol approved by the Institutional Animal Care and Use Committee at UC Davis.

### Sex as a Biological Variable

Both male and female mice were included in all experiments in the present study.

### Preparation of single cell suspensions from WT and TAZ KO mice

Six corneas from 3 mice (2 female and 1 male) with the same genotype (WT or TAZ KO) and age (2 or 11 months) were dissected and pooled together in 600 µL of papain solution (Papain Vial; CAT LK003178; Worthington Biochemical Corp., Lakewood, NJ) and incubated in a warm water bath at 37 °C for 1.75 hours. The tube was gently inverted every 10-20 minutes during incubation to improve cell dissociation. The cell suspension in papain solution was aspirated and passed through the Flowmi cell strainer (40 µM; CAT 136800040, Bel-Art, Wayne, NJ) and the strainer was flushed with 300 µL of 1X PBS to collect additional cells. The filtered cell suspension was centrifuged at 400 g for 10 min at 4 °C and the supernatant was discarded. The cell pellet was resuspended on 100 µL of 0.04% bovine serum albumin (BSA) in 1X PBS and centrifuged at 300 g for 5 min at 4 °C, which was repeated twice. The final pellet was resuspended on 40 µL of 2% BSA-PBS. Single cell suspensions were prepared in the same manner for each genotype and age group. All single cell suspensions were submitted to the UC Davis DNA Technologies and Expression Analysis Core for 10X Chromium Single Cell 3’ library preparation and NovaSeq 4000 sequencing.

### Single cell RNA-sequencing analysis

Raw FASTQ sequencing data was processed by the UC Davis Bioinformatics Core using 10X CellRanger for alignment to the 10x Genomics mouse reference genome (*GRCh38-mm10*). Further analysis was then performed on R using *Seurat* v5 (45). Count matrices for each age were converted to Seurat objects. The objects were independently normalized, scaled and reduced using the function *Seurat::SCTransform* v2 (*glmGamPoi*) with cell cycle regression and the unfiltered object was subject to dimensionality reduction (determined using knee identification in elbow plot), k-means clustering (resolution determined using *clustree* package) and neighbor identification (46–48). Low quality clusters were identified and filtering based on low gene counts, high mitochondrial read percentage and high expression of MHC class I genes (49). Ambient RNA decontamination was performed on individual objects using the *DecontX* package as described in the package documentation (50). The objects were re-clustered and cells with a high ambient fraction were removed. Objects were integrated over age using the *Harmony* package (16). The embeddings were utilized for Uniform Manifold Approximation Projection (UMAP) and subsequent clustering and neighbor identification as described prior.

Cell type assignment was performed using marker genes defined in murine single cell literature as well as using top markers for each cluster identified by *Seurat::FindAllMarkers* (Wilcoxon-Rank-Sum) (17,51,52). Canonical marker genes were identified from cross-species corneal transcriptional literature (17,18,53–55).

This *Seurat::FindAllMarkers* was also performed for differentially expressed gene (DEG) identification across samples where DEGs were defined by *P < 0.05* and *|logFC| > 0*. To further minimize the impact of ambient RNA contamination on DEG analysis, we eliminated genes with a high contribution to the contamination fraction using package *DropletUtils* (56). In brief, *DropletUtils::ambientProfileEmpty* was used to define the ambient RNA profile from the raw count matrix for each sample.

*DropletUtils::ambientContribMaximum* was then used to provide a contamination fraction for each gene within each sample. The sum of contamination fractions for each gene in both samples of comparison for the DEG analysis was utilized to filter highly contaminant genes (contamination fraction *<0.1*). The filtered DEG list was utilized for visualization and further analysis via over-representation analysis with gene ontology (GO ORA) (package *clusterprofiler*) (57). An intersection approach was performed to identify shared pathways between ages in TAZ KO mice in order to avoid over-representation of samples. Genes upregulated in the TAZ KO CEnC independent of age were determined by the intersection of genes upregulated in TAZ KO CEnC between 2-and 11-months of age (compared to the WT CEnCs). Genes downregulated in the TAZ KO CEnC independent of age were determined by the intersection of genes downregulated in TAZ KO CEnC between 2- and 11-months of age (compared to the WT CEnCs).

To assess gene pathways (percentage mitochondrial gene and ER Stress), *Seurat::AddModuleScore* was utilized to calculate select pathway scores using the genes from gene ontology (GO) terms with the mouse genome annotation (obtained from package *org.Mm.eg.db*). Non-normality of the data was tested via Shapiro-Wilks test (*P* < 2.2e-16 for percent mitochondrial gene score and *P* < 0.001 for ER stress score) and visualization via histogram in **Supplementary File 1 S1**. Thus, Wilcoxon rank sum testing was applied to determine statistical differences in scores between groups.

Figures were generated using Seurat plotting functions accompanied by ggplot2, ggpubr, and rstatix packages (58–60). Figures were organized using BioRender.

### Transmission electron microscopy (TEM) of CEnCs

Mice were euthanized, and enucleated eyes were immediately fixed in 2.5% glutaraldehyde and 2% paraformaldehyde in 0.1 M sodium cacodylate buffer. After 30 minutes, corneas were dissected and then fixed in 2.5% glutaraldehyde and 2% paraformaldehyde (PFA) in 0.1 M sodium cacodylate buffer again. Three corneas of each age group of WT and TAZ KO mice were dissected. Electron microscopy of the cornea was performed with a transmission electron microscope (TEM; Talos L120C with Ceta CMOS digital camera (4 K × 4 K), F.E.I. Company, Hillsboro, OR) as previously described (61).

The number of mitochondria was counted in 3 TEM images taken from each cornea at X36,000 magnification. Mitochondria with indistinct cristae and electrolucent areas were deemed abnormal, and the ratio of the number of dysfunctional mitochondria/the total number of mitochondria was calculated for each image and averaged for each mouse (62). 2-way ANOVA was used to compare across the genotypes. *P* < 0.05 was considered statistically significant.

### Immunofluorescence staining in CEnC wholemounts

Immunofluorescence staining was performed on the corneal endothelium as previously described (15). For ZO-1 (1:100, Invitrogen, #61-7300) and Na,K-ATPase (ATP1A1) staining (1:200, Invitrogen, MA5-32184), images from our previous publication were reviewed again for qualitative assessment of the morphology and distribution of each staining between genotypes (15).

For GRP78 staining, a primary antibody against GRP78 (1:50, Proteintech, 11587-1-AP) was used. Seven corneas were collected from WT and 9 corneas were collected from TAZ KO mice and stained as previously described. To quantitatively compare the staining intensity of GRP78, corrected total cell fluorescence (CTCF) was calculated using the ImageJ software, according to previously described protocols to control for local background fluorescence and the following formula was used: CTCF = integrated density − (area of selected cell × mean fluorescence of background readings) (63). Unpaired t-test was used to compare between genotypes; *P*<0.05 was considered statistically significant. For ATP synthase staining, a primary antibody against ATP synthase (1:200, Sigma Aldrich, A9728) was used. Eight corneas from WT and 8 corneas from TAZ KO mice were stained. The staining intensity of ATP synthase was analyzed by calculating and comparing CTCF as described above.

## Data Availability

Single cell transcriptomic data for the wildtype mouse corneas are available on NCBI GEO (GSE267704). TAZ KO data will be available with the same accession upon publication.

## Acknowledgements and funding sources

This study was supported by the National Institutes of Health (NIH) R01EY016134, R01EY036440, R01EY037135, P30EY12576, K08EY028199, and T32EY015387.

## Commercial Relationships Disclosure

S. Park, None; R. Ramarapu, None; J. Lim, None; S. Khan, None; M. Khan, None; W. Stoehr, None; B.C. Leonard, None; S.M. Thomasy, None

