## Supplementary file 1 for "TAZ (*Wwtr1*) deficiency leads to ER stress and mitochondrial dysfunction in a mouse model of Fuchs’ endothelial corneal dystrophy"

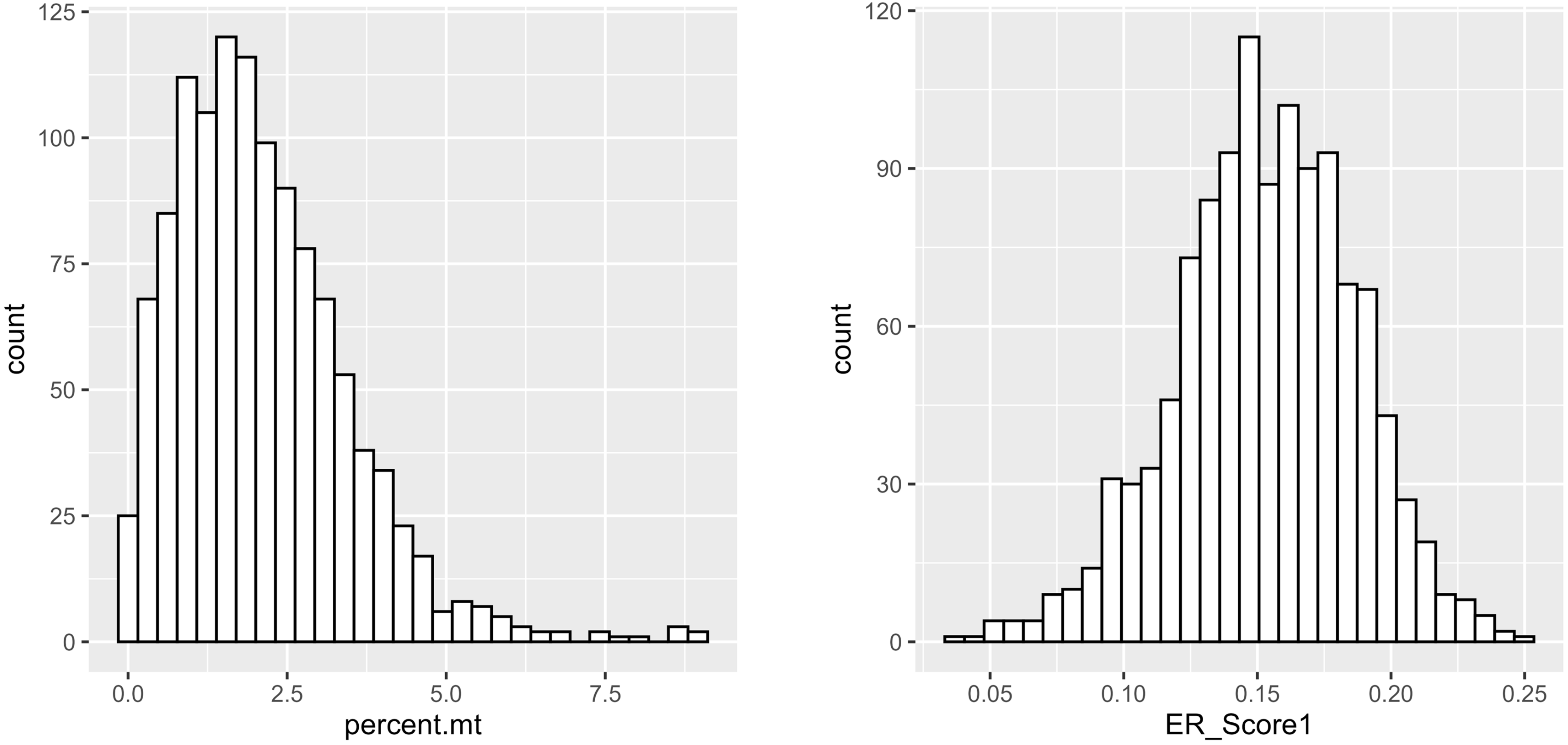


**S1. Data distribution of gene scores across CEnC.** (A) histogram of percentage mitochondrial gene score. (B) Histogram of ER stress gene score.

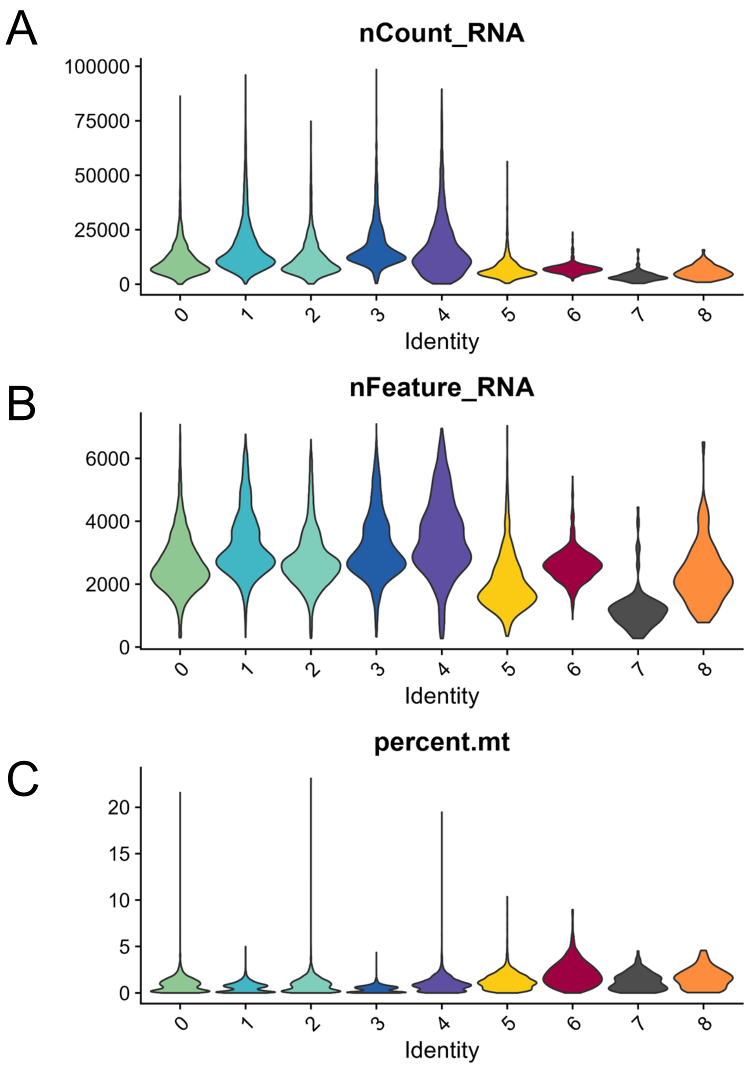


**S2. Quality control plots for single cell RNA-sequencing.** (A) Violin plot of number of transcripts per cell by cluster. (B) Violin plot of number of unique transcripts per cell by cluster. (C) Violin plot of average percentage of mitochondrial transcripts by cluster.

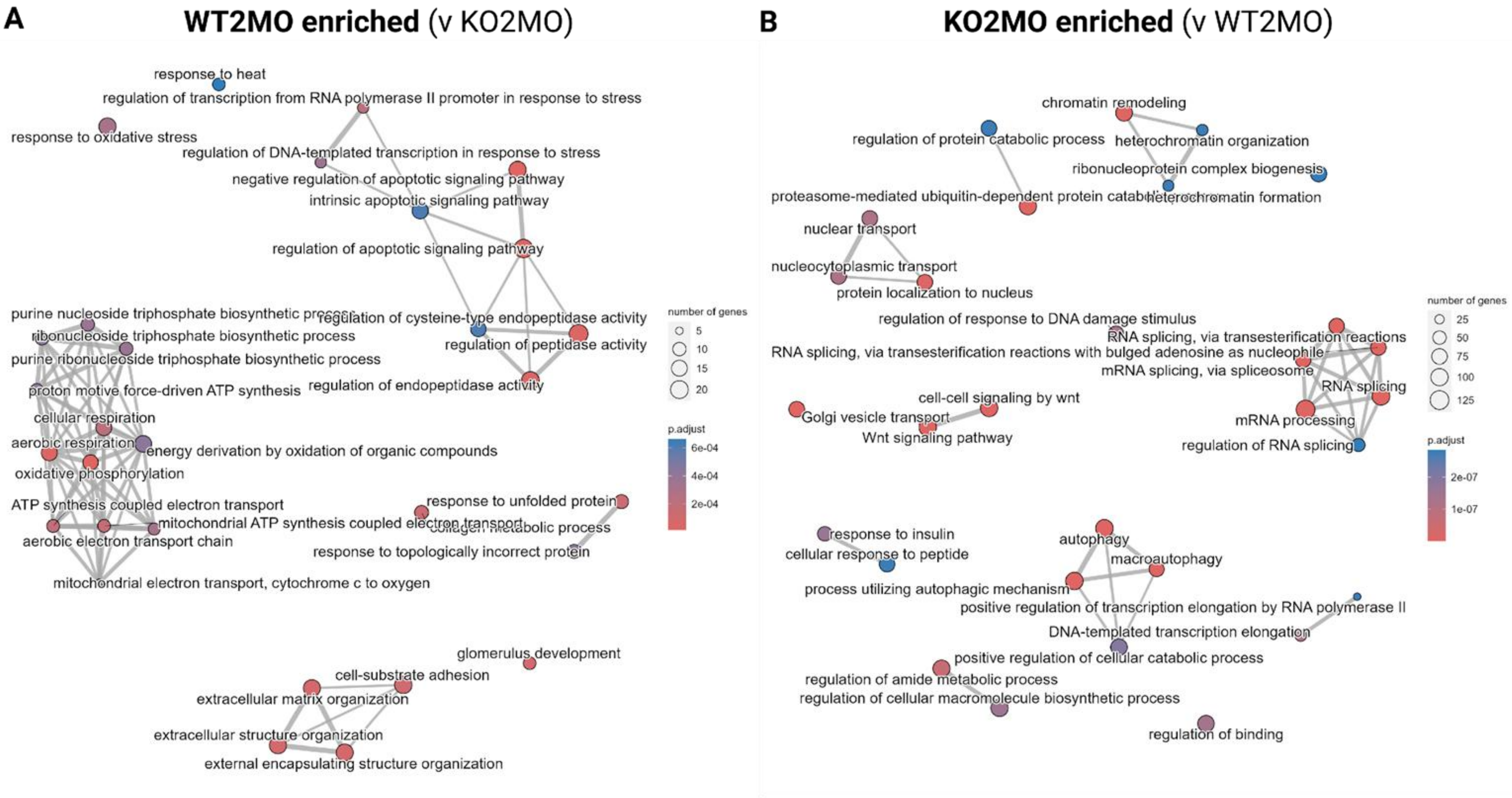


**S3. GO ORA for DEG analysis between WT and TAZ KO at 2 months of age** (A) Net plot of GO biological process terms enriched in WT compared to TAZ KO at 2 months of age. (B) Net plot of GO biological process terms enriched in TAZ KO compared to WT at 2 months of age.

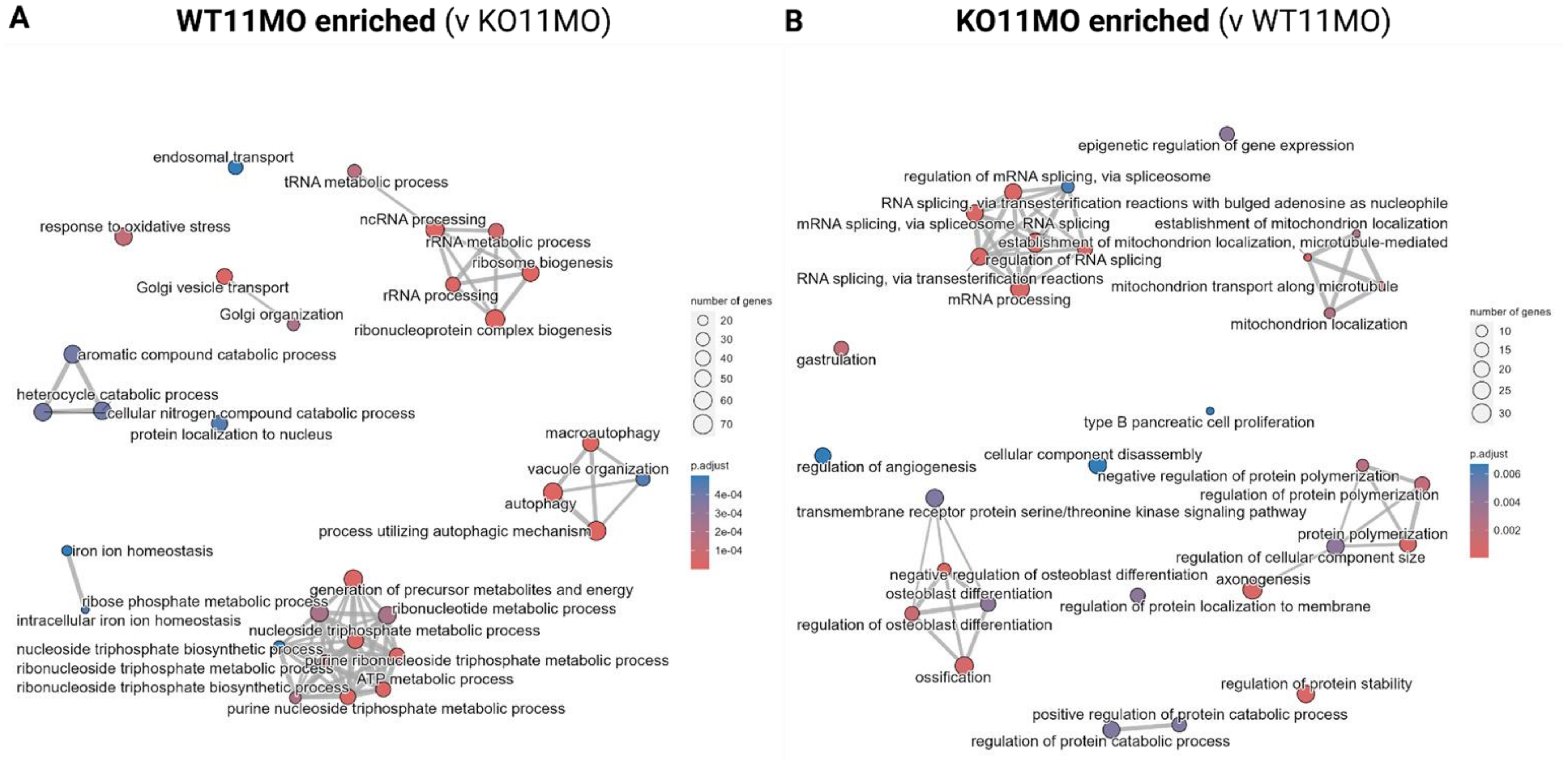


**S4. GO ORA for DEG analysis between WT and TAZ KO at 11 months of age** (A) Net plot of GO biological process terms enriched in WT compared to TAZ KO at 11 months of age. (B) Net plot of GO biological process terms enriched in TAZ KO compared to WT at 11 months of age.

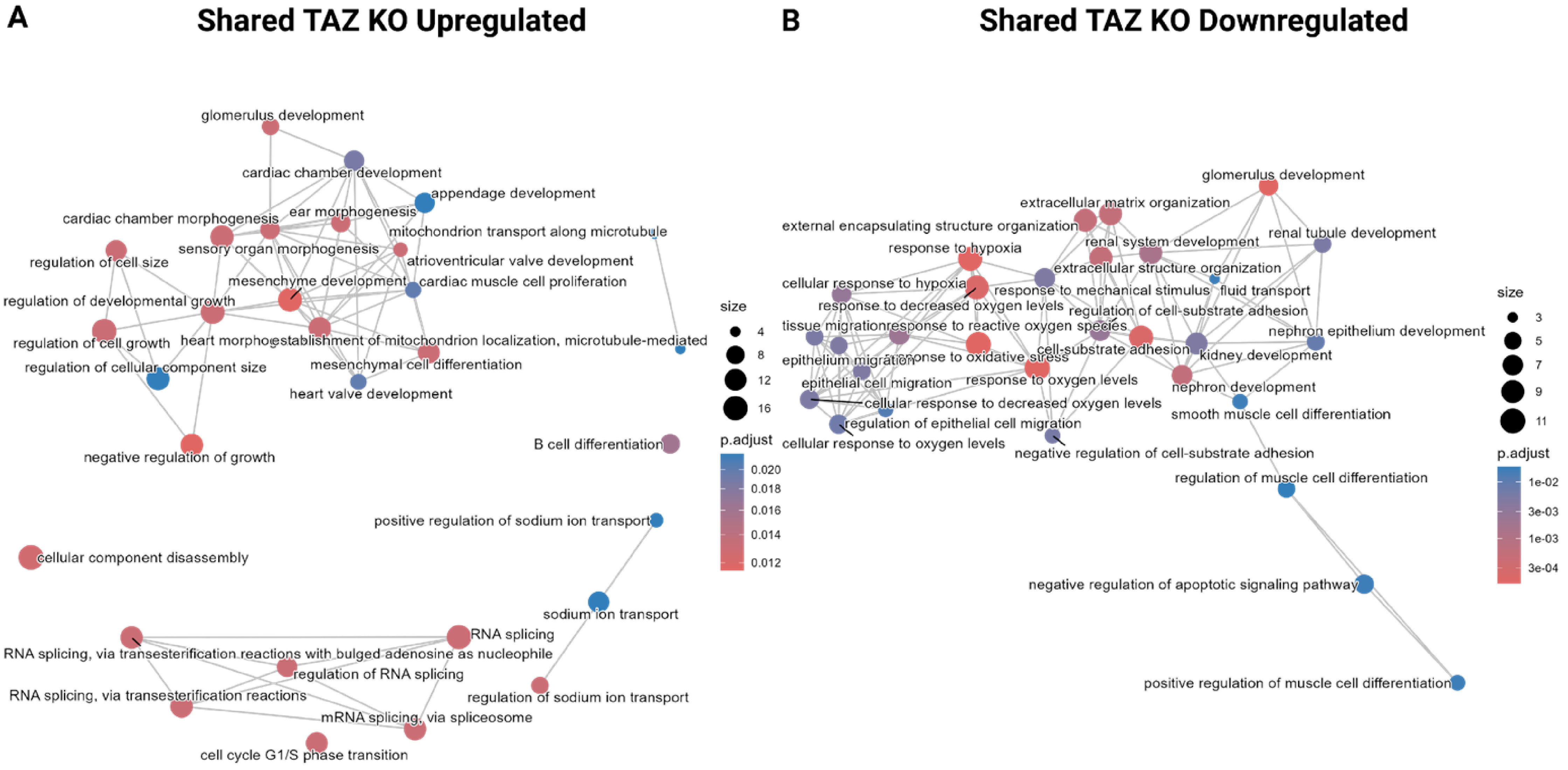


**S5. GO ORA for Shared TAZ KO upregulated and downregulated gene sets** (A) Net plot of GO biological process terms enriched in TAZ KO CEnC of age. (B) Net plot of GO biological process terms depleted in TAZ KO CEnC of age.

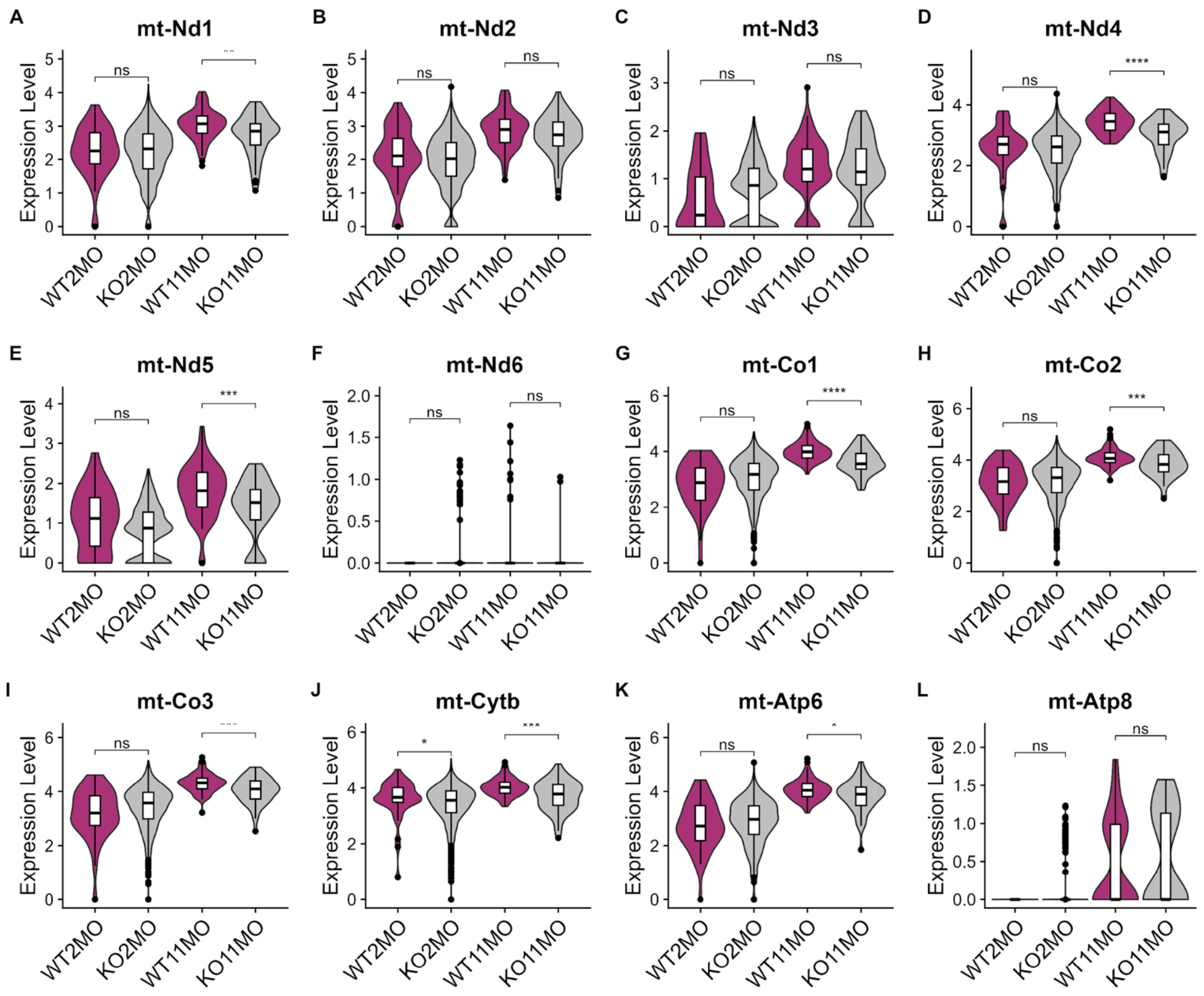


**S6. Expression of mitochondrial genes across CEnC samples** (A-I) Violin plots of mitochondrial genes.
